# Very low amplitude muscle activity increases probability of motor evoked potentials in healthy individuals and in ALS

**DOI:** 10.1101/2025.08.07.663223

**Authors:** Séamus O’Sullivan, Yasmine Tadjine, Narin Suleyman, Eva Woods, Antonio Fasano, Cathal Walsh, Friedemann Awiszus, Orla Hardiman, Richard G. Carson, Roisin McMackin

## Abstract

**Introduction:** In many clinical and research settings, transcranial magnetic stimulation (TMS) intensities are standardised based on resting motor threshold (RMT). It is well-established that contraction of the target muscle increases motor evoked potential (MEP) amplitude and correspondingly decreases RMT. As such, when estimating RMT it is crucial to ensure the target muscle is relaxed. Typically trials in which baseline electromyographic (EMG) amplitude exceeds a specified threshold are rejected. The influence of motor activity below typical rejection thresholds on MEP amplitudes has yet to be established.

**Methods:** We retrospectively analysed TMS-EMG data collected during RMT measurement in 45 healthy controls (1761 datapoints) and 35 people with amyotrophic lateral sclerosis (ALS, 1238 datapoints). Trials where root mean squared (RMS) baseline EMG amplitude exceeded 10*µV* were rejected. Generalised linear mixed-effects models were used to assess effects of muscle activity below this rejection threshold on probability of evoking an MEP with peak-to-peak amplitude ≥50*µV*.

**Results:** Greater sub-rejection-threshold activity significantly increases MEP probability in controls (p<0.0004) and people with ALS (p=0.0010). Models predicted a 31-38% increase in MEP probability when baseline RMS-EMG amplitude increased from 1*µV* to 9*µV*. Sub-rejection– threshold baseline activity was significantly greater in ALS than controls (p=0.0055).

**Discussion:** We have shown for the first time that higher EMG amplitudes below a typical rejection threshold markedly increase probability of evoking an MEP with peak-to-peak amplitude ≥50*µV*. Researchers should take measures to account for effects of sub-rejection–threshold activity on RMT, particularly in populations where baseline activity may be elevated, such as in ALS.

## 1. Introduction

Transcranial magnetic stimulation (TMS) is frequently employed to study cortical and corticospinal function and infer that there are changes in cortical excitability in various neurological and psychiatric disorders [1]. In addition, repetitive TMS is often used for therapeutic purposes [2]. In both research and clinical settings, the strength of the magnetic field that constitutes an adequate intensity of stimulation is typically standardized on the basis of each individual’s resting motor threshold (RMT). Classically, this is defined as the minimum stimulation intensity required to evoke a motor evoked potential (MEP) of at least 50µV peak-to-peak amplitude in a target muscle, in 50% of trials (i.e. with a probability of 0.5) [3].

It is well established that voluntary contraction of the target muscle increases MEP amplitude [4], decreases onset latency [5] and reduces amplitude variability [6]. This has been demonstrated at low levels of muscle force, such as 5% of maximum voluntary contraction (MVC) [7]. Furthermore, muscle contraction increases the slope of stimulus-response curves [8] and reduces the stimulation intensity at which a motor response is first evoked [4]. To ensure comparability across study populations or experimental conditions, it is therefore considered essential that the RMT, and other resting MEP-based measures, are obtained while the motoneurons innervating the target muscle are not in receipt of excitatory drive. To achieve this objective, protocols used to determine the RMT (and other “resting” measures) typically reject MEPs recorded when a target muscle shows pre-stimulus EMG activity.

When the specific threshold used for MEP rejection is reported, it is typically a root mean squared EMG amplitude (RMS-EMG) in the target muscle over 2.5*µV* [9], 5*µV* [10,11], 10*µV* or 20*µV* [12,13], during a brief (e.g., 100 ms) period immediately prior to stimulation. On occasion however, much higher thresholds of 50*µV* [14] or 100*µV* [15,16] RMS-EMG have been employed. To date, however, it has not been established whether variation in background RMS-EMG below cut-off thresholds typically employed (e.g., 10*µV*) influences the probability that an MEP will be evoked. Any such influence would significantly impact the inferences that can be drawn when comparing RMTs and related “resting” MEP measures between groups or conditions that differ in respect of background EMG. To exemplify this point, we consider studies that engage individuals living with amyotrophic lateral sclerosis (ALS).

ALS is a neurodegenerative disorder involving the degeneration of upper and lower motoneurons. As it has a wide range of presentations, and is prone to misdiagnosis [17], reliable biomarkers for early detection are urgently required. It has been proposed that TMS-based measures have the potential to meet this need [18]. Differences between people with ALS and controls in respect of both single-pulse (e.g., RMT) and paired-pulse TMS measures (e.g., short intracortical inhibition, SICI) suggest that those with ALS exhibit cortical hyperexcitability [19–21]. Abnormalities in SICI appear to be present even in individuals with few clinical upper motor neuron signs [20]. As such, it is now recommended that SICI be used as a supportive diagnostic biomarker of ALS [22]. It is, however, widely appreciated that individuals with ALS show elevated resting EMG activity (relative to controls) and additional distinguishing features such as fasciculations [23]. Estimates of SICI are based on either differences in MEP amplitude (conditioned versus unconditioned, for fixed intensity protocols) or the probability of evoking MEPs of specific amplitude (for threshold tracking protocols). Since the stimulation intensities employed in both protocols are defined relative to RMT, factors that exert a systematic influence on estimation of the RMT (such as the elevated resting EMG activity characteristic of ALS), have the potential to affect measures such as SICI.

In the present study, we aimed to test the hypothesis that variation in background EMG activity below a typical rejection threshold (RMS-EMG *<* 10*µV*) is systematically related to the probability of evoking an MEP of peak-to-peak amplitude ≥ 50*µV*. This was accomplished through a retrospective analysis of data collected during the tracking of RMT in both healthy controls and people with ALS.

## 2. Methods

### 2.1. Ethical Approval

Ethical approval was obtained from the ethics committee of St. James’s Hospital (REC reference: 2017-02). All participants were over 18 years of age and provided written consent prior to participation. With the exception of preregistration, all work was performed in accordance with the Declaration of Helsinki.

### 2.2 Participants

Control participants were recruited from the community. All participants with ALS were diagnosed with Possible, Probable or Definite ALS according to the revised El Escorial Criteria [24]. People with ALS were recruited from the Irish National ALS Clinic, Beaumont Hospital, Dublin, Ireland. Participants were excluded if they had TMS contraindications as per the revised TMS screening questionnaire [25], or a neuromuscular or neurodegenerative disorder other than ALS. Only right-handed participants were included, i.e., those with a laterality quotient of zero or higher on the Edinburgh Handedness Inventory [26]. Participants with a resting motor threshold (RMT) greater than 99 percent of maximum stimulator output (%MSO) were excluded.

### 2.3 Experimental Protocol

#### 2.3.1 Electromyography

Participants were seated comfortably in a sofa-style chair with wide armrests. Their arms were positioned at an angle of 90−120° on the armrests or in the participant’s lap to improve comfort and promote muscle relaxation. Ag-AgCl electrodes (Cleartrance 1700, Conmed, Haverhill, MA, USA) were placed on the abductor pollicis brevis (APB), spaced approximately 2cm apart in a belly-tendon montage. A reference electrode was placed on the ulnar styloid of the right wrist. Bipolar EMG was recorded from the right hand. The signal was amplified, with gain 1000, and band-pass filtered at 10-1000Hz using BioPac EMG100C amplifiers (Biopac Systems UK, Pershore, UK). Ambient electrical noise was removed, either using a Humbug Noise Eliminator or equivalently, a D400 Multichannel Noise Eliminator (Digitimer Ltd., Welwyn Garden City, UK). The signal was digitised at 10kHz using a Cambridge Electronic Design Micro1401 digitiser (CED, Cambridge, UK) and recorded using Signal software version 7.0.1 (CED, Cambridge, UK).

#### 2.3.2 Transcranial Magnetic Stimulation

Monophasic magnetic pulses were delivered across the scalp using a DuoMag MP Dual stimulator (Deymed Diagnostics, s.r.o., Hronovo, Czech Republic) equipped with a 50mm mid-diameter figure-of-eight coil. The axis of intersection between the two loops of the coil was angled 45° relative to the saggital plane, inducing posteroanterior (PA) current across the primary motor cortex (M1). The TMS “hotspot”, defined as the point on the scalp requiring the lowest stimulation intensity (SI) to evoke MEPs in the APB, was identified. To ensure consistent coil placement, 7 landmarks corresponding to markers on the surface of the coil were marked on a cloth cap securely taped to the participant’s head (outlined in detail in McMackin et al., 2024).

This study included TMS-EMG data collected as part of a prior study of differences in TMS-based measures in ALS compared to controls. Single-pulse TMS was delivered as part of an automated threshold-tracking protocol using Maximum Likelihood Parameter Estimation by Sequential Testing (PEST). The protocol, developed by Prof. Friedemann Awiszus [27], uses a sigmoid-shaped logistic function to determine the %MSO at which there is a 50% probability of eliciting an MEP with a peak-to-peak amplitude above a specific target. In the case of the data retrospectively included in this study, this target amplitude was either 50*µV* (i.e., while recording RMT) or 200*µV* (i.e., while recording the “threshold hunting target”) as described in McMackin et al. (2024).

A fully automatic implementation of the PEST algorithm, in MATLAB R2016a (MathWorks Inc., MA, USA) and Signal Software (CED Ltd., Cambridge, UK), was employed to reduce the likelihood of human error and enable online rejection of trials with excessive baseline EMG activity, as described in detail in McMackin et al. (2024). In all cases, where RMS-EMG in the 200ms immediately prior to stimulation exceeded 10*µV*, data were not passed to the PEST algorithm and a replacement trial was performed.

### 2.4 Data Analysis

#### 2.4.1 Models

The effect of RMS-EMG on MEP probability was assessed using generalised linear mixed effects models, as implemented in lme4 [28]. Modelling was performed separately for people with ALS and controls as a suitable model convergence could not be achieved when employing a single model with an additional “diagnosis” factor. All statistical analysis was performed in R version 4.4.0 [29].

Logistic models had a categorical response variable of MEP/no MEP, which was equal to one if an MEP with a peak-to-peak amplitude of ≥ 50*µV* was evoked, or zero otherwise. As the data were collected as part of a prior threshold-tracking study, stimulus intensity was not fixed within or across individuals. Correspondingly, the normalised SI (%MSO as a percentage of RMT) applied in each trial was accounted for as part of the modelling process. A factor of “Participant” was also included in all models to account for inter-individual variability. Akaike Information Criterion (AIC) and Bayesian Information Criterion (BIC) were used to compare competing models with nuisance parameters of age and sex, and fixed effects of RMS-EMG and normalised SI as well as the interaction between these two factors. These parameters were included in the final analysis if their inclusion resulted in a lower AIC and BIC in either ALS or control models. All continuous numeric parameters were z-transformed to prevent model convergence issues.

It was found that including a random intercept frequently resulted in a singular fit, and had a low variance. The model intercept was therefore not allowed to vary randomly by subject. This is equivalent to assuming that all participants had a similar probability of evoking an MEP with a peak-to-peak amplitude ≥50*µV* when all predictors are set to zero (i.e., their mean value, because all predictors were z-transformed during model fitting). The low variance of the intercept is likely caused by the normalisation of stimulus intensity to each participant’s RMT. By definition, RMT is the SI at which there is a 50% probability of evoking an MEP with peak-to-peak amplitude ≥50*µV*. Therefore, at a fixed value of SI, MEP probability is likely to be similar across participants.

Model significance was assessed by comparison to a null model, containing a parameter only for stimulation intensity, normalised to RMT (i.e., given as a percentage of RMT). Parametric bootstrapping, employing 10,000 simulations determined by the R package pbkrtest version 0.5.2 [30] was used to produce p-values for model comparisons. Parametric bootstrapping works by using the null model to generate predicted values, creating a simulated data set resembling that which might be observed under the null hypothesis. A null model and full model (containing a parameter for RMS-EMG) are then fit to the simulated data, and a likelihood ratio calculated. This procedure is repeated a large number of times to generate a null distribution for the likelihood ratio test statistic. This distribution can then be used to calculate a p-value for the likelihood ratio obtained from models fit to the observed data. P-values are calculated by dividing the number of simulations in which the simulated likelihood ratio was more extreme than the observed likelihood ratio by the total number of simulations, i.e., the “PBtest” method from pkbrtest. For more detail on this procedure, refer to Halekoh and Højsgaard (2014). Predicted values of MEP probability for each model (see Figure 1) were generated using the package ggeffects version 1.7.0 [31].

**Figure 1.**
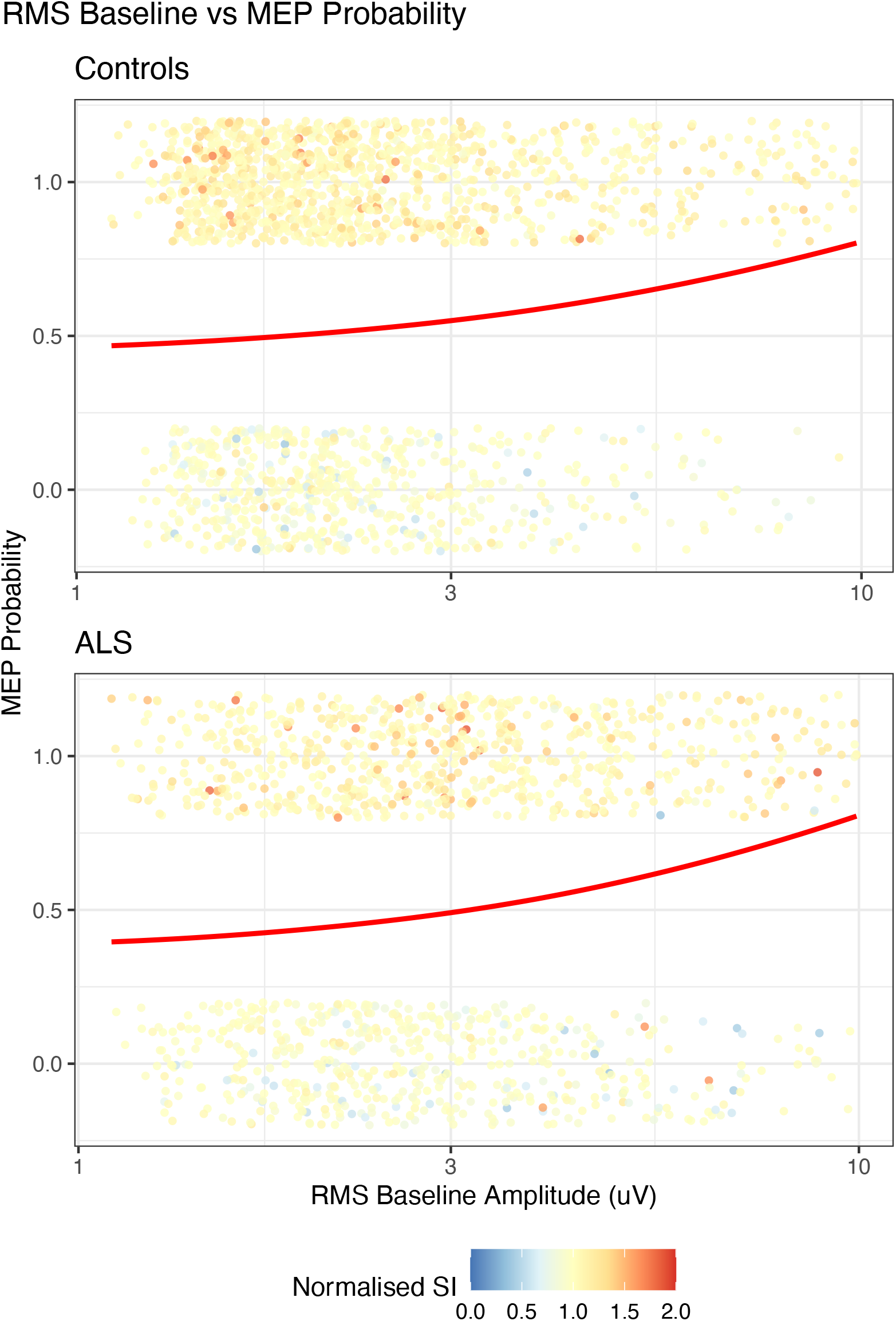
Predicted values for MEP probability, when normalised SI is 100% RMT, across RMS-EMG values (x-axis, shown on log scale). Red line represents the model-predicted probability of evoking an MEP with a peak-to-peak amplitude. Coloured points represent individual TMS trials, in which a motor evoked potential (MEP) either occurred (top) or did not occur (bottom). Points are jittered to avoid overlap. SI – Stimulation intensity, RMS – Root mean square

#### 2.4.2 RMS-EMG Group Comparison

Median RMS-EMG amplitudes of participants with ALS and controls were compared with an unpaired, two-tailed Mann-Whitney U Test, using the built-in wilcox.test function in R 4.4.0 [29].

## 3. Results

### 3.1 Included Cohort

Data from 45 controls (1761 TMS-EMG trials in total) and 35 people with ALS (1238 TMS-EMG trials in total) were included in this study. Details of these cohorts are summarized in table Table 1. Thirty participants with ALS in this study were concurrently taking riluzole, which is found to have no significant effect on RMT [32,33].

**Table 1.**
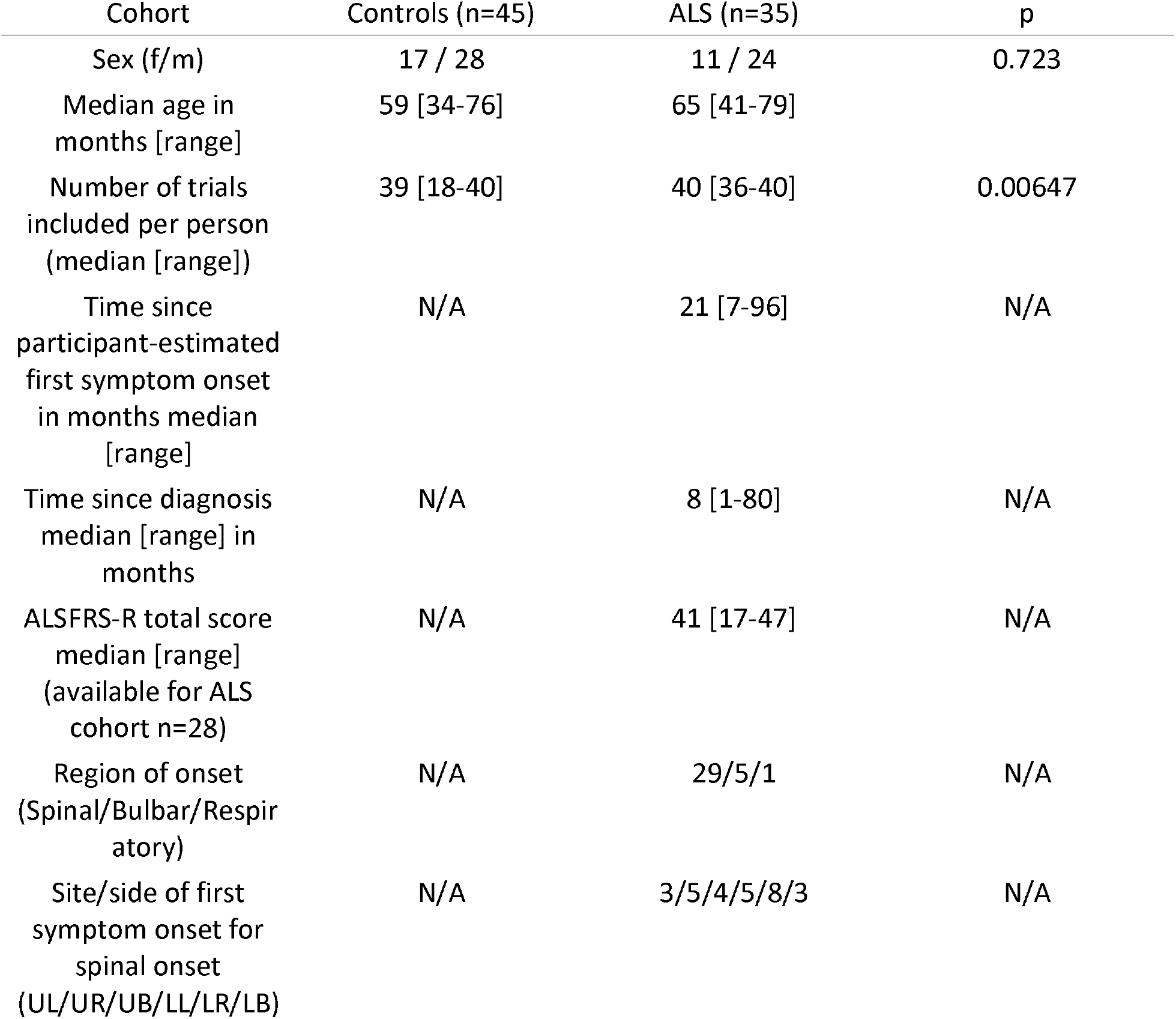
Summary of statistics for ALS and control cohorts. P-value for sex was determined by x^2^ testing. Age and number of trials p-values were determined by sign rank testing. ALSFRS-R – Revised ALS functional rating scale. UL – Upper left limb. UR – Upper right limb. UB – Both upper limbs. LL – Lower left limb. LR – Lower right limb. LB – Both lower limbs.

| Cohort | Controls (n=45) | ALS (n=35) | p |
| --- | --- | --- | --- |
| Sex (f/m) | 17 / 28 | 11 / 24 | 0.723 |
| Median age in months [range] | 59 [34-76] | 65 [41-79] |  |
| Number of trials included per person (median [range]) | 39 [18-40] | 40 [36-40] | 0.00647 |
| Time since participant-estimated first symptom onset in months median [range] | N/A | 21 [7-96] | N/A |
| Time since diagnosis median [range] in months | N/A | 8 [1-80] | N/A |
| ALSFRS-R total score median [range] (available for ALS cohort n=28) | N/A | 41 [17-47] | N/A |
| Region of onset (Spinal/Bulbar/Respiratory) | N/A | 29/5/1 | N/A |
| Site/side of first symptom onset for spinal onset (UL/UR/UB/LL/LR/LB) | N/A | 3/5/4/5/8/3 | N/A |

### 3.2 Model Selection and Comparison

The following model formula was chosen based on optimisation of AIC and BIC:

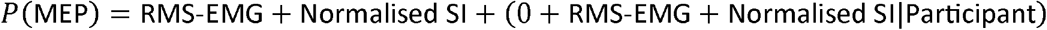

Notably, inclusion of the RMS-EMG term improved model fit in both people with ALS and controls.

Results of the model comparison are displayed in Table 2.

**Table 2.** Results of model-comparison-based hypothesis test comparing models with and without a term for RMS-EMG. Parametric bootstrapping with 10,000 simulations was used to obtain p-values for the model comparison. See methods for more detail.

| Group | Likelihood Ratio | P-value |
| --- | --- | --- |
| Controls | 16.374 | 0.0004 |
| ALS | 22.344 | 0.0010 |

Predicted values for the probability of obtaining an MEP with a peak-to-peak amplitude of ≥50*µV*, while stimulus intensity is fixed at 100% RMT, are shown for a sample of sub-threshold RMS-EMG amplitudes in Figure 1 and Table 3.

**Table 3.** Predicted values for MEP probability when normalised stimulus intensity is 100% RMT, across three different values of RMS-EMG. RMS – Root mean square

| RMS-EMG ($\mu V$) | Controls | ALS |
| --- | --- | --- |
| 1 | 0.463 | 0.390 |
| 5 | 0.633 | 0.593 |
| 9 | 0.775 | 0.769 |

### 3.3 RMS-EMG Group Comparison

The result of the Mann-Whitney U test comparing the average RMS-EMG activation between groups is shown in Figure 2. Median RMS-EMG values were significantly higher in people with ALS than in controls (p=0.0055).

**Figure 2.**
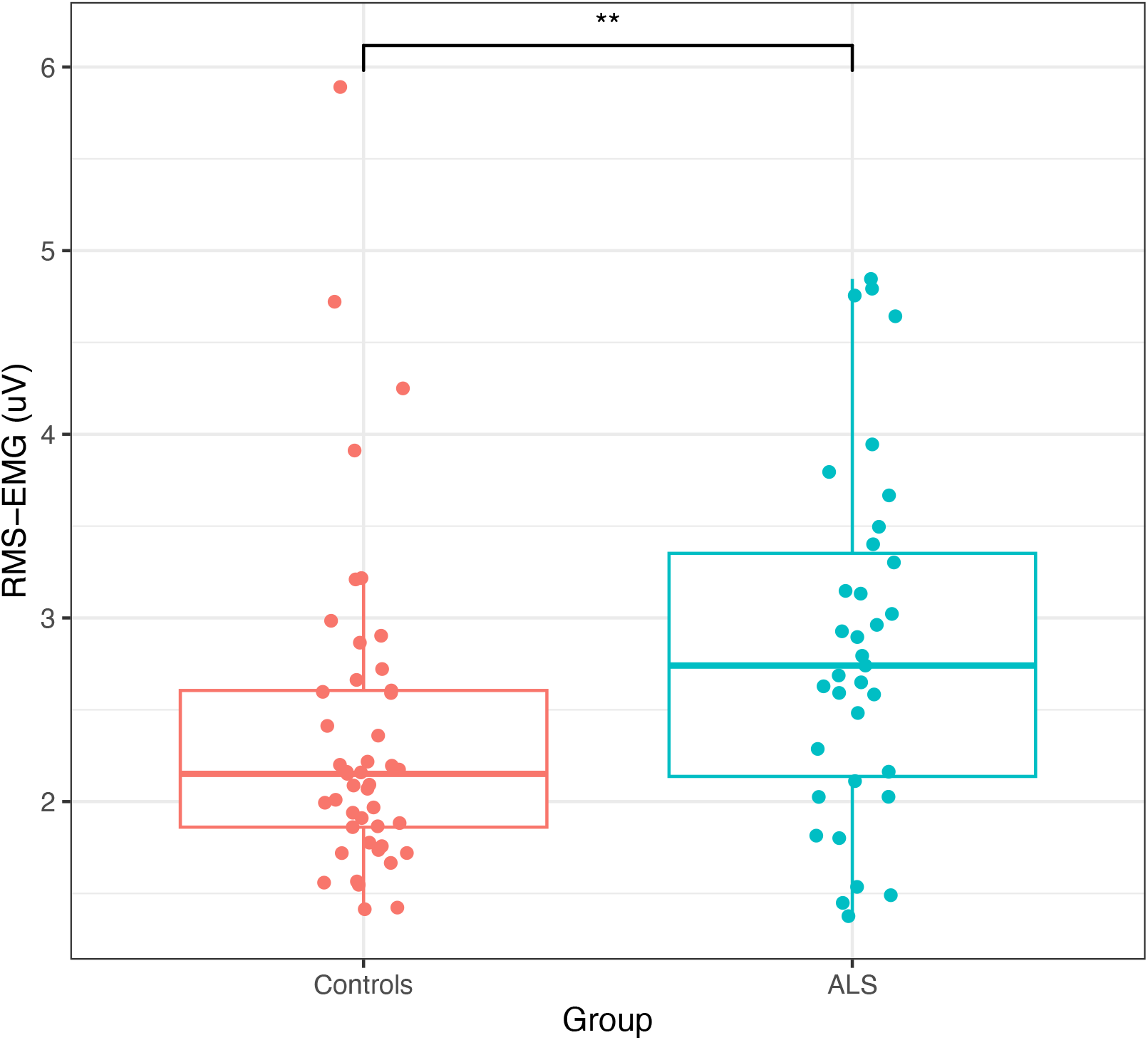
Comparison of median RMS-EMG amplitudes in controls and people with ALS across all trials. Boxplots display mean and interquartile range (IQR). Whiskers extend to the largest value no greater than 1.5 times the IQR. (** p < 0.01, * p < 0.05, Mann-Whitney U Test). RMS – Root mean square. ALS – Amyotrophic lateral sclerosis

## 4 Discussion

In this study we have demonstrated that, within the range 0 − 10*µV*, variations in background EMG RMS amplitude systematically influence the probability of evoking an MEP of at least 50*µV*. Moreover, the size of this effect is non-negligible – in healthy controls an increase in RMS EMG activity from 1*µV* to 9*µV* results in more than a 31% increase in the probability of evoking an MEP that meets the standard 50*µV* amplitude criterion. An increase from 1*µV* to 5*µV* corresponded to an increase in probability of 17%. We found that this effect could be replicated in data from a cohort of people with ALS, with increase in RMS EMG activity from 1*µV* to 9*µV* resulting in a 38% increase in MEP probability and an increase from 1*µV* to 5*µV* corresponded to a 20% increase in MEP probability. Thus, our findings indicate that estimated RMT values are likely to vary inversely with background muscle activity, even when trials with RMS-EMG ≥ 10*µV* are excluded.

To our knowledge, no other study has explicitly modelled the effects of variations in RMS-EMG below 10*µV*. The findings have important implications for research which employs TMS to study motor networks in health and disease. Currently, there is no standard procedure to control for the presence of background muscle activity. Baseline EMG amplitude rejection thresholds vary significantly from study to study, and can range from 2.5*µV* [9] to 100*µV* [15,16]. In addition, many studies do not report a rejection threshold, or the time window within which background EMG is monitored [34]. Several investigators seek to verify relaxation simply by online visual or auditory monitoring of the EMG.

These findings have important implications for the definition and recording of resting motor threshold. RMT is defined as the TMS stimulus intensity for which there is a 50% probability of evoking an MEP at rest. However, an important assumption of threshold estimation, including via the PEST-based approach employed to estimate RMT here, is that MEP probability is constant for a given stimulus strength. Our modelling shows that even for the range of RMS-EMG values within which a muscle is defined as “resting”, there is significant variation in MEP probability. This is supported by previous findings of state-dependent changes in RMT, e.g., during motor imagery [35]. Therefore, the RMT measures we have employed to normalize stimulation intensity across individuals in our model may not reflect motor threshold at complete rest.

It is particularly important to consider the potential effects of EMG activity that is below the nominal rejection threshold when a study involves people with neurological disorders, such as ALS, that affect motoneuron excitability. We have demonstrated that in a sample of people with ALS, background EMG in trials which would typically be deemed acceptable (i.e. background RMS-EMG no greater than 10*µV*) is significantly greater than in controls.

Consequently, investigators should assess carefully how the effects of such differences in background muscle activity can be mitigated when employing outcome measures based on MEP amplitudes. Our modelling (see Figure 1) suggests that the effect of variations in background muscle activity on MEP amplitudes is likely to be small for levels of RMS-EMG below 2 *µV* (the slope of the curve shown in the figure is negligible for such values). In practice however, the adoption of this stringent criterion (particularly for people with neurological disorders) will likely lead to extensive rejection and repetition of trials, thus prolonging the required duration of recording sessions. This can often be a disincentive to participation, and induce fatigue or further muscle tension during recording. An alternative would be to include RMS-EMG as a covariate in statistical models, with the aim of accounting for the influence of its variation on outcome measures. Alternatively, studies employing protocols dependent upon RMT could continuously monitor RMT and suitably account for fluctuations in the degree of “rest” during recording. Such an approach has been adopted in some threshold tracking TMS studies of ALS[20,36], as well as a recent fixed-intensity TMS study in controls[37]. In these studies, the estimated RMT, and conditioning stimulus intensities dependent thereon, are continuously adjusted to account for fluctuations in cortical excitability [38]. At a minimum, in all TMS-EMG-based studies, investigators should clearly state (i) the time window in which pre-stimulation muscle activity is registered, (ii) the specific threshold of baseline EMG activity above which trials will be rejected, and (iii) the presence of any differences in EMG activity between experimental conditions or cohorts. In this regard, inferential tests of equivalence should be used.

## 6. Acknowledgements

We thank the Wellcome-HRB Clinical Research Facility at St. James’s Hospital for providing a dedicated environment for the conduct of high-quality clinical research. Finally, we would like to thank all the people with ALS, participants and their families who volunteered to take part in this study.

